# Hyaluronidase: the spreading factor of *Tityus serrulatus* venom

**DOI:** 10.1101/487298

**Authors:** Bárbara Bruna Ribeiro de Oliveira-Mendes, Sued Eustáquio Mendes Miranda, Douglas Ferreira Sales-Medina, Bárbara de Freitas Magalhães, Yan Kalapothakis, Renan Pedra de Souza, Valbert Nascimento Cardoso, André Luís Branco de Barros, Clara Guerra-Duarte, Evanguedes Kalapothakis, Carolina Campolina Rebello Horta

## Abstract

**Background:** The hyaluronidase enzyme is generally known as a spreading factor in animal venoms. Although its activity has been demonstrated in several organisms, a deeper knowledge about hyaluronidase and the venom spreading process from the bite/sting site until its elimination from the victim's body is still in need.

**Methods and principal findings:** We used technetium-99m radiolabeled *Tityus serrulatus* venom (^99m^Tc-TsV) to evaluate the venom distribution kinetics in mice. To understand the hyaluronidase’s role in the venom’s biodistribution, ^99m^Tc-TsV was immunoneutralized with specific anti-*T.serrulatus* hyaluronidase serum. Venom biodistribution was monitored by scintigraphic images of treated animals and by measuring radioactivity levels in tissues as heart, liver, lungs, spleen, thyroid, and kidneys. In general, results revealed that hyaluronidase inhibition delays venom components distribution, when compared to the non-neutralized ^99m^Tc-TsV control group. Scintigraphic images showed that the majority of the immunoneutralized venom is retained at the injection site, whereas non-treated venom is quickly biodistributed throughout the animal’s body. At the first 30 minutes, concentration peaks are observed in the heart, liver, lungs, spleen, and thyroid, which gradually decreases over time. On the other hand, immunoneutralized ^99m^Tc-TsV takes 240 minutes to reach high concentrations in the organs. A higher concentration of immunoneutralized ^99m^Tc-TsV was observed in the kidneys in comparison with the non-treated venom. Further, *in situ* neutralization of ^99m^Tc-TsV by anti-*T.serrulatus* hyaluronidase serum at zero, ten, and 30 minutes post venom injection showed that late inhibition of hyaluronidase can still affect venom biodistribution. In this assay, immunoneutralized ^99m^Tc-TsV was accumulated in the bloodstream until 120 or 240 minutes after TsV injection, depending on anti-hyaluronidase administration time. Altogether, our data show that immunoneutralization of hyaluronidase prevents venom spreading from the injection site.

**Conclusions:** The results obtained in the present work show that hyaluronidase has a key role not only in the venom spreading from the inoculation point to the bloodstream, but also in venom biodistribution from the bloodstream to target organs. Our findings demonstrate that hyaluronidase is indeed an important spreading factor of TsV, and its inhibition can be used as a novel first-aid strategy in envenoming.

**Author summary:** Hyaluronidases are known as the venom components responsible for disseminating toxins from the injection site to the victim’s organism. Therefore, understanding how the venom distribution occurs and the role of hyaluronidases in this process is crucial in the field of toxinology. In this study, we inhibited *Tityus serrulatus* venom (TsV) hyaluronidase’s action using specific anti-Ts-hyaluronidase antibodies. Labeling TsV with a radioactive compound enabled monitoring of its biodistribution in mice. Our results show that, upon hyaluronidase inhibition, TsV remains at the injection site for longer, and only a reduced amount of the venom reaches the bloodstream. Consequently, the venom arrives later at target organs like the heart, liver, lungs, spleen, and thyroid. Considering the possible application of hyaluronidase inhibition as a therapeutic resource in envenoming first-aid treatment, we performed the administration of hyaluronidase neutralizing antibodies at different times after TsV injection. We observed that TsV remains in the bloodstream and its arrival at tissues is delayed by 120 or 240 minutes after TsV injection, depending on anti-hyaluronidase administration times. Our data show that hyaluronidase plays a crucial role in TsV spreading from the injection site to the bloodstream and from the bloodstream to the organs, thus suggesting that its inhibition can help to improve envenoming’s treatment.

## Introduction

Scorpionism is considered a serious public health threat and was officially recognized as a neglected tropical disease by the Brazilian Academy of Sciences [1]. In Brazil, scorpion sting reports have been increasing over the years, reaching 90,000 accidents in 2016, and outnumbering the accidents caused by other venomous animals such as spiders and snakes [2].

The yellow scorpion *Tityus serrulatus* (Ts) (Lutz and Mello Campos, 1922) is considered the most venomous scorpion in South America [3–5], causing serious envenomation accidents mainly in southeast Brazil [6] and representing the species of greatest medical-scientific importance in the country.

The symptomatology of scorpionism involves local pain, which can be associated with nausea, sweating, tachycardia, fever, and stirring. Moderate complications may include epigastric pain, cramps, vomiting, hypotension, diarrhea, bradycardia, and dyspnea. Severe envenoming may present several potentially lethal complications, such as cardio-respiratory failure [7–11].

These symptoms are related to the synergic action of a variety of toxic components present in the venom. Ts venom (TsV) consists of a complex mixture of components such as mucus, lipids, amines, nucleotides, inorganic salts, hyaluronidases, serine proteases, metalloproteases, natriuretic peptides, bradykinin potentiating peptides, antimicrobial peptides, high molecular weight (Mw) proteins, and ion channel active neurotoxins, which are the major toxic components [12–24].

Hyaluronidases are extensively found in the venoms of various animals such as snakes, scorpions, spiders, and others [25]. Venom hyaluronidases are always referred to as "spreading factors" [26,27], as they hydrolyze the hyaluronic acid (HA) present in the interstitial matrix, thus helping the venom toxins to reach the victim’s bloodstream and invade its organism. Hyaluronidase’s enzymatic action increases membrane absorbency, reduces viscosity, and makes tissues more permeable to injected fluids (spreading effect). Therefore, hyaluronidase acts as a catalyst for systemic envenoming [25].

TsV hyaluronidase activity was first demonstrated by Possani’s group [28], and the enzyme was later isolated and partially characterized by Pessini and collaborators [14]. Horta et al. [16] further expanded these studies by performing extensive molecular, biological, and immunological characterization of TsV hyaluronidase. The authors described the sequence of two enzyme isoforms showing 83% identity, Ts-Hyal-1 and Ts-Hyal-2, by cDNA analysis of the venom gland. A purified native Ts hyaluronidase was used to produce anti-hyaluronidase serum in rabbits. Epitopes common to both isoforms were mapped, and it was shown that they include active site residues. Most importantly, it was demonstrated for the first time that *in vivo* neutralization assays with anti-hyaluronidase serum inhibited and delayed mouse death after injection of a lethal dose of TsV, thus confirming the influence of hyaluronidase in TsV lethality [16].

The active recombinant hyaluronidase Ts-Hyal-1 from TsV was produced and characterized. It is an important biotechnological tool for the attainment of sufficient amounts of the enzyme for structural and functional studies [29].

Herein, we further pursued the goal of demonstrating the importance of hyaluronidase as a spreading factor of TsV. Our results show that inhibition of the hyaluronidase activity of TsV in mice hinders venom spreading from the injection site as well as its biodistribution to the tissues.

## Methods

### Scorpions and venom extraction

*T. serrulatus* scorpions were collected in Belo Horizonte, Minas Gerais, Brazil, with proper licensing from the competent authorities (IBAMA, Instituto Brasileiro do Meio Ambiente e dos Recursos Naturais Renováveis, protocol number 31800-1). Venom was obtained from female scorpions regularly milked twice a month by electrical stimulation of telson. After extraction, venom was solubilized in ultrapure water and centrifuged at 16,000g at 4°C for 10 min. The supernatant was quantified using Bio-Rad “Protein DC assay” kit [30], and stored at -20°C until use.

### Experimental animals

Female Swiss CF1 mice (6-8 weeks old, 24-28 g) were obtained from the animal care facilities (CEBIO) of the Federal University of Minas Gerais (UFMG). Animals had free access to water and food and were kept under controlled environmental conditions.

### Ethics statement

The Ethics Committee (Comissão de Ética no Uso de Animais, CEUA) of UFMG certifies that the procedures using animals in this work are in agreement with the Ethical Principals established by the Brazilian Council for the Control of Animal Experimentation (CONCEA). Protocol number 05/2016. Approved: March 8, 2016.

### Anti-hyaluronidase serum

The anti-hyaluronidase serum used in this work was produced by Horta et al. [16] through immunization of rabbits with native hyaluronidase purified from TsV.

### Hyaluronidase activity: *In vitro* neutralization assay

Hyaluronidase activity was measured according to the turbidimetric method described by Pukrittayakamee et al. [31] with modifications [16]. The assay mixture contained 12.5 µg of HA (Sigma-Aldrich), acetate buffer (0.2 M sodium acetate-acetic acid pH 6.0, 0.15 M NaCl), and test (or control) sample in a final volume of 250 µl. Commercial hyaluronidase from bovine testis (12.5 µg; Apsen) was used as a positive control, and ultra-pure water was used as a negative control. Assay mixtures were incubated for 15 min at 37°C, and reactions were stopped by adding 500 µl of stop solution containing 2.5% (w/v) cetyltrimethylammonium bromide (CTAB) dissolved in 2% (w/v) NaOH. Assays were monitored by absorbance at 400 nm against a blank of acetate buffer (250 µl) and stop solution (500 µl). Turbidity of the samples decreased proportionally to the enzymatic activity of hyaluronidase. Values were expressed as percentages of hyaluronidase activity relative to the negative (no addition of enzyme, 0% activity) and positive (addition of commercial enzyme, 100% activity) controls.

The tested samples were TsV (2 µg) and TsV neutralized with anti-hyaluronidase serum (2 µg of TsV incubated for 1 h at 37°C with 10 µl of anti-hyaluronidase serum).

### Radiolabeling of *T. serrulatus* venom (TsV)

To label TsV with technetium-99m (^99m^Tc; IPEN São Paulo), a sealed vial containing 200 μg of SnCl_2_.H_2_O solution in 0.25 mol/l HCl (2 mg/ml) and 50 μg of NaBH_4_ solution in 0.1 mol/l NaOH (1 mg/ml) was prepared. The pH was adjusted to 7.4 using 1 mol/l NaOH. Next, 25 μl of TsV (5 g/l in saline 0.9% w/v) was added, and vacuum was applied to the vial, followed by addition of 0.1 ml of Na^99m^TcO^4^ (3.7 MBq). The solution was kept at room temperature for 15 min.

### Radiochemical purity evaluation

Radiochemical purity was determined by thin layer chromatography (TLC-SG, Merck) using acetone as the mobile phase to quantify ^99m^TcO_4_ -. Strips radioactivity was determined by a gamma counter (Wallac Wizard 1470-020 Gamma Counter, PerkinElmer Inc.). ^99m^TcO_2_ was removed from the preparation using a 0.45 μm syringe filter [32].

### *In vitro* stability of ^99m^Tc-TsV

Tests in saline 0.9% (w/v) and in mice plasma were performed to evaluate the stability of the radiolabeled complex ^99m^Tc-TsV.

TLC-SG was used to evaluate the stability of the radiolabeled complex diluted in saline. The labeled solution was kept at room temperature, and aliquots were taken at 1, 2, 4, 6 and 24 h for determination of radiochemical purity.

A volume of 90µl ^99m^Tc-TsV solution was incubated with 1 ml of fresh mouse plasma at 37°C under agitation. Radiochemical stability was determined by TLC-SG from samples taken at 1, 2, 4, 6 and 24 h after incubation.

### Blood clearance of ^99m^Tc-TsV

An amount equivalent to 3.7 MBq of ^99m^Tc-TsV was diluted (10% v/v) in phosphate-buffered saline (PBS; control) or anti-hyaluronidase serum and incubated for 1h at 37°C. Then, 50 µl of the samples were intramuscularly injected into the right tight of healthy Swiss mice (6-8 weeks old, 24-28 g; n = 6 per group). Mice were anesthetized with a mixture of xylazine (15 mg/kg) and ketamine (80 mg/kg), and an incision was made in the animals’ tails for blood collection in pre-weighed tubes at 1, 5, 10, 15, 20, 30, 45, 60, 90, 120, 240, and 1440 minutes after administration of the samples. The tubes were weighted, and their radioactivity determined by a gamma counter. These data were used to plot the percentage of dose injected per gram tissue (% ID/g) versus time.

### Scintigraphic images of mice injected with ^99m^Tc-TsV

Aliquots of 18 MBq of ^99m^Tc-TsV in 10% (v/v) PBS (control) or anti-hyaluronidase serum (pre-incubated for 1h at 37°C) were intramuscularly injected (50 µl) into the right tight of healthy Swiss mice (6-8 weeks old, 24-28 g; n = 3 per group). Animals were anesthetized at 30, 60, and 120 min after sample administration with a mixture of ketamine (80 mg/kg) and xylazine (15 mg/kg) and placed horizontally under a gamma camera (Nuclide TM TH 22, Mediso). Images were collected with a Low Energy High Resolution (LEHR) collimator and 256×256×16 dimension matrices with a 300 s acquisition time, using a 20% symmetrical window with a fixed energy peak at 140 KeV.

### Biodistribution of ^99m^Tc-TsV

Aliquots of 3.7 MBq of ^99m^Tc-TsV in 10% (v/v) PBS (control) or anti-hyaluronidase serum (pre-incubated for 1h at 37°C) were intramuscularly injected (50 µl) into the right tight of healthy Swiss mice (6-8 weeks old, 24-28 g; n = 6 per group). Mice were euthanized at 30, 60, 240, and 1440 min post-injection, and heart, liver, lungs, spleen, thyroid, and kidneys were dissected, dried with filter paper, and weighed. The radioactivity in each tissue was determined by a gamma counter. A standard dose containing the same injected amount of ^99m^Tc-TsV was counted simultaneously in a separate tube, which was defined as 100% radioactivity. The results were expressed as the percentage of injected dose per gram of tissue (%ID/g).

### Evaluation of hyaluronidase neutralization as a first-aid treatment

An amount equivalent to 3.7 MBq of ^99m^Tc-TsV diluted in PBS was intramuscularly injected (25 µl) into the right tight of healthy Swiss mice (6-8 weeks old, 24-28 g; n = 6 per group). Next, anti-hyaluronidase serum was inoculated (25 µl, intramuscularly) into the same site of ^99m^Tc-TsV injection at different time-points (0, 10, and 30 min post-injection of ^99m^Tc-TsV). Mice were anesthetized with a mixture of xylazine (15 mg/kg) and ketamine (80 mg/kg), and an incision was made in the animals’ tails for blood collection in pre-weighed tubes at 1, 5, 10, 15, 20, 30, 45, 60, 90, 120, and 240 minutes after administration of ^99m^Tc-TsV. The tubes were weighted, and their radioactivity determined by a gamma counter. Data were used to plot the percentage of dose injected per gram tissue (% ID/g) versus time.

### Statistical analyses

Sample sizes were calculated using G Power version 3.1. To compare multiple means, the sample size was calculated considering alpha (α), power effect, effect size (f), and population size (n). To estimate number of mice needed in the ^99m^Tc-TsV biodistribution assays, parameters were set at f = 0.7, α = 0.05, power = 0.8, and groups = 4. Data were expressed as mean ± S.E.M. Graphs were plotted using the software GraphPad PRISM version 5.00 (La Jolla, CA, USA).

All statistical tests were carried out on R version 3.4.4. Significance level was set at 0.05, and tests were performed two-sided. Effect of serum administration and time on the mean ^99m^Tc-TsV biodistribution was evaluated using two-way ANOVA. Normality and equal variance suppositions were assessed using Shapiro-Wilk and Levene’s tests, respectively. Effects of serum administration and time on ^99m^Tc-TsV mean blood clearance were analyzed using a linear mixed model (lme function on nlme package).

## Results

### Anti-hyaluronidase serum neutralizes TsV hyaluronidase activity *in vitro*

In the turbidimetric assays, commercial hyaluronidase from bovine testis exhibited high hyaluronidase activity, which was referred to as 100% activity (positive control). Ultra-pure water had no enzyme activity, which was referred to as 0% activity (negative control). TsV presented high enzymatic activity, similar to that observed for the commercial hyaluronidase (positive control) and was considered to be 100% active (Fig. 1). In the *in vitro* neutralization assay, pre-incubation of TsV with anti-hyaluronidase serum completely neutralized hyaluronidase activity (Fig. 1).

**Figure 1.**
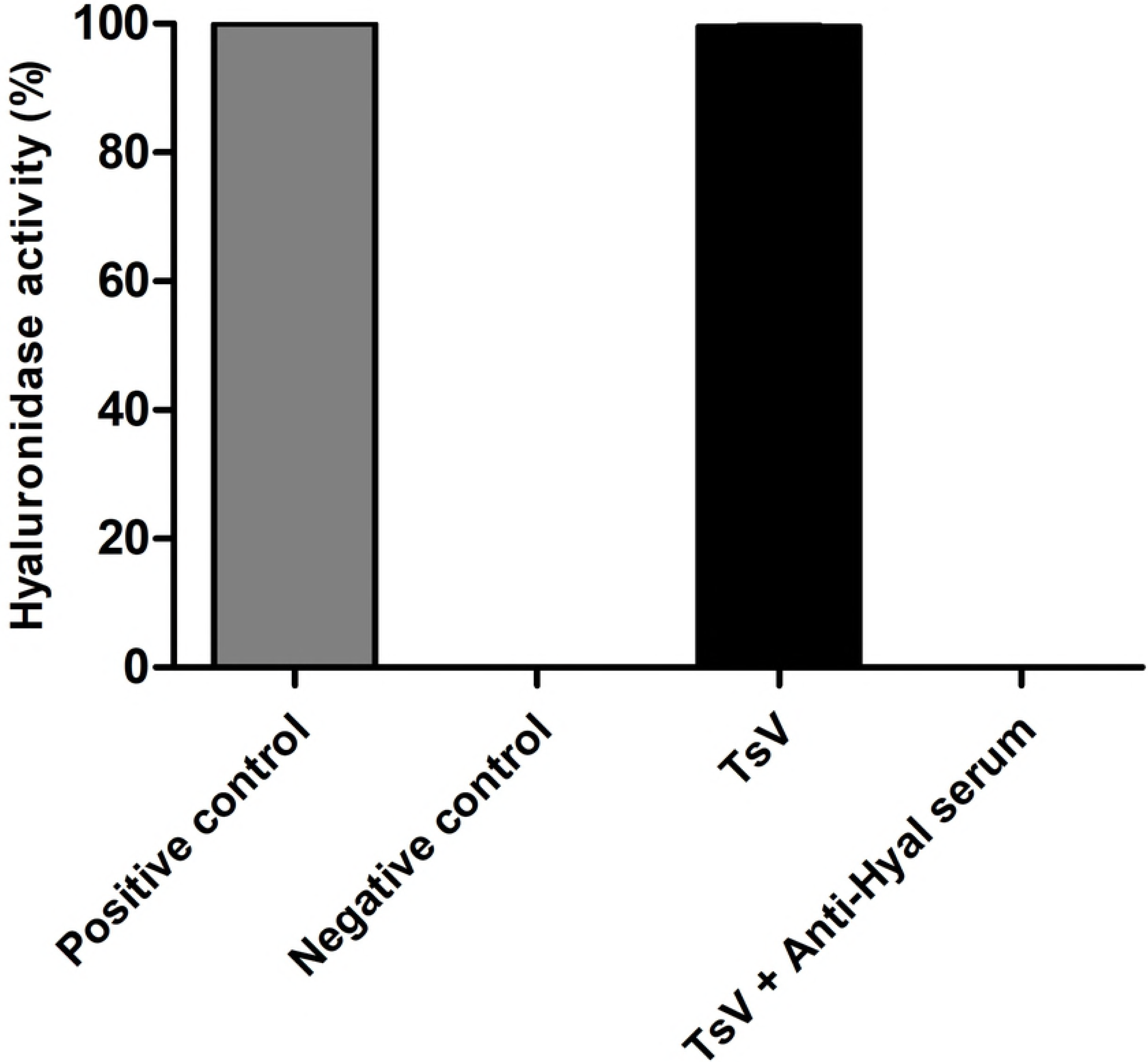
*In vitro* neutralization assay using rabbit anti-hyaluronidase serum. Hyaluronidase activity (%) was measured using a turbidimetric assay. Commercial hyaluronidase from bovine testis was used as a positive control, and ultrapure water was used as a negative control. TsV (2 µg) was incubated with anti-hyaluronidase serum (Anti-Hyal, 10 µl) for 1 h at 37°C before testing. Anti-hyaluronidase serum neutralized 100% of the hyaluronidase activity in TsV. All values are expressed as the mean ± S.E.M. of duplicates from three independent experiments.

### ^99m^Tc-TsV presents high radiochemical yields and radiolabeling stability

The radiochemical efficiency of the TsV labeling with technetium-99m was determined by TLC. The results indicated high radiochemical yield (95.2 ± 2.4%; data not shown). Radiochemical yields higher than 90% are recommended for *in vivo* application of radiopharmaceuticals [33]. Therefore, our ^99m^Tc-TsV complex presented suitable radiochemical characteristics, which encouraged further *in vivo* studies.

The radiolabeling stability curve for ^99m^Tc-TsV is shown in Fig. 2. Stability tests were performed after 1, 2, 4, 6 and 24 h of incubation of ^99m^Tc-TsV in saline 0.9% (w/v) at room temperature or in fresh mouse plasma at 37°C. High stability was observed over long periods of time (>95%), thus indicating suitability for further biodistribution assays.

**Figure 2.**
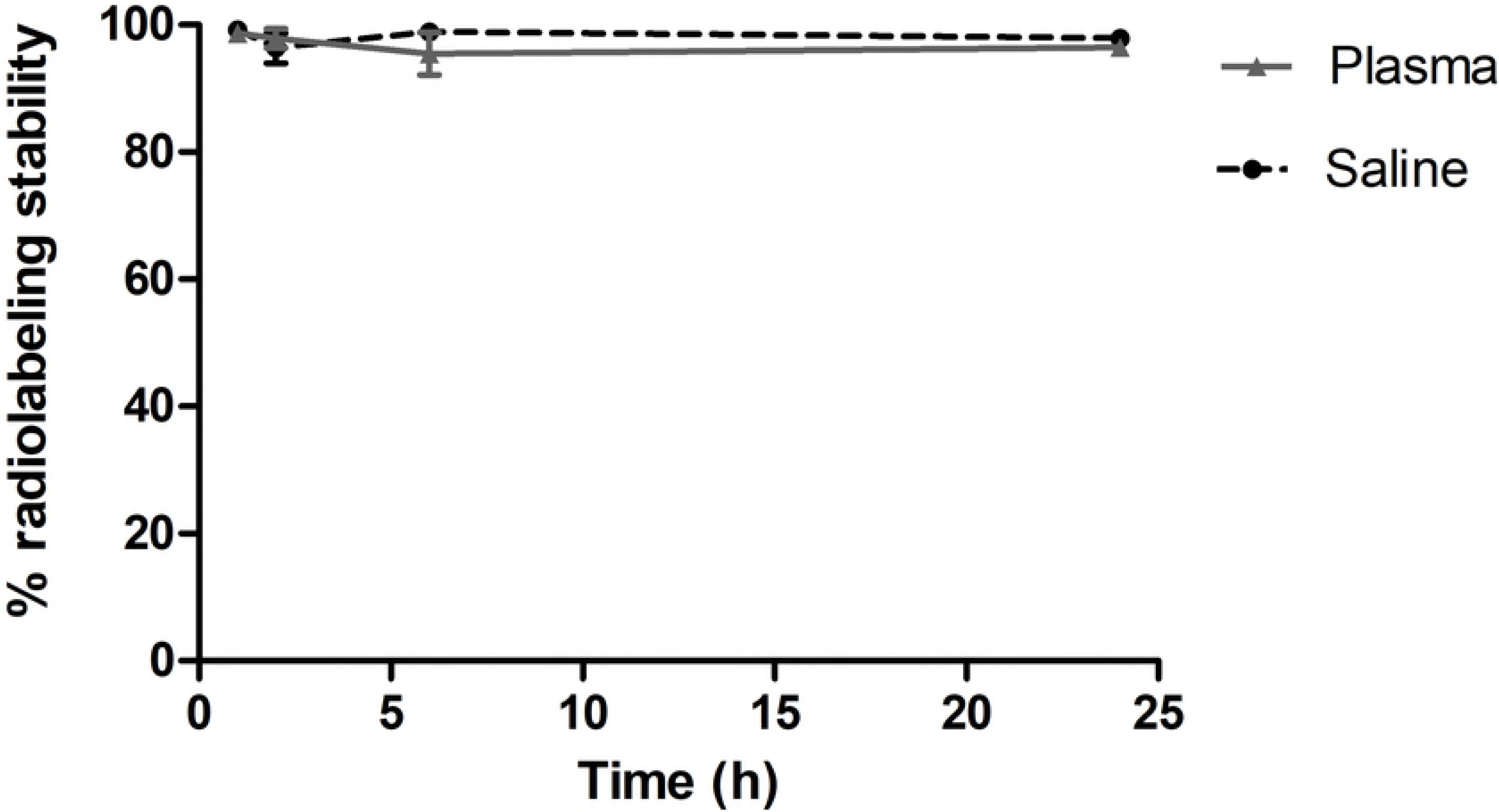
*In vitro* stability of ^99m^Tc-TsV. Stability of the complex ^99m^Tc-TsV over time in the presence of saline 0.9% (w/v) at room temperature and in the presence of plasma at 37°C. All values are presented as the mean ± S.E.M. of duplicates from three independent experiments.

### Neutralization of hyaluronidase impairs TsV spreading

Blood clearance of ^99m^Tc-TsV diluted in PBS or pre-neutralized with anti-hyaluronidase serum is shown in Fig. 3. Following the injection into healthy Swiss mice, the ^99m^Tc-TsV complex showed quick absorption, reaching the highest bloodstream levels after 30 min. After this time point, ^99m^Tc-TsV concentration in the bloodstream decreases, which indicates biodistribution to the tissues.

**Figure 3.**
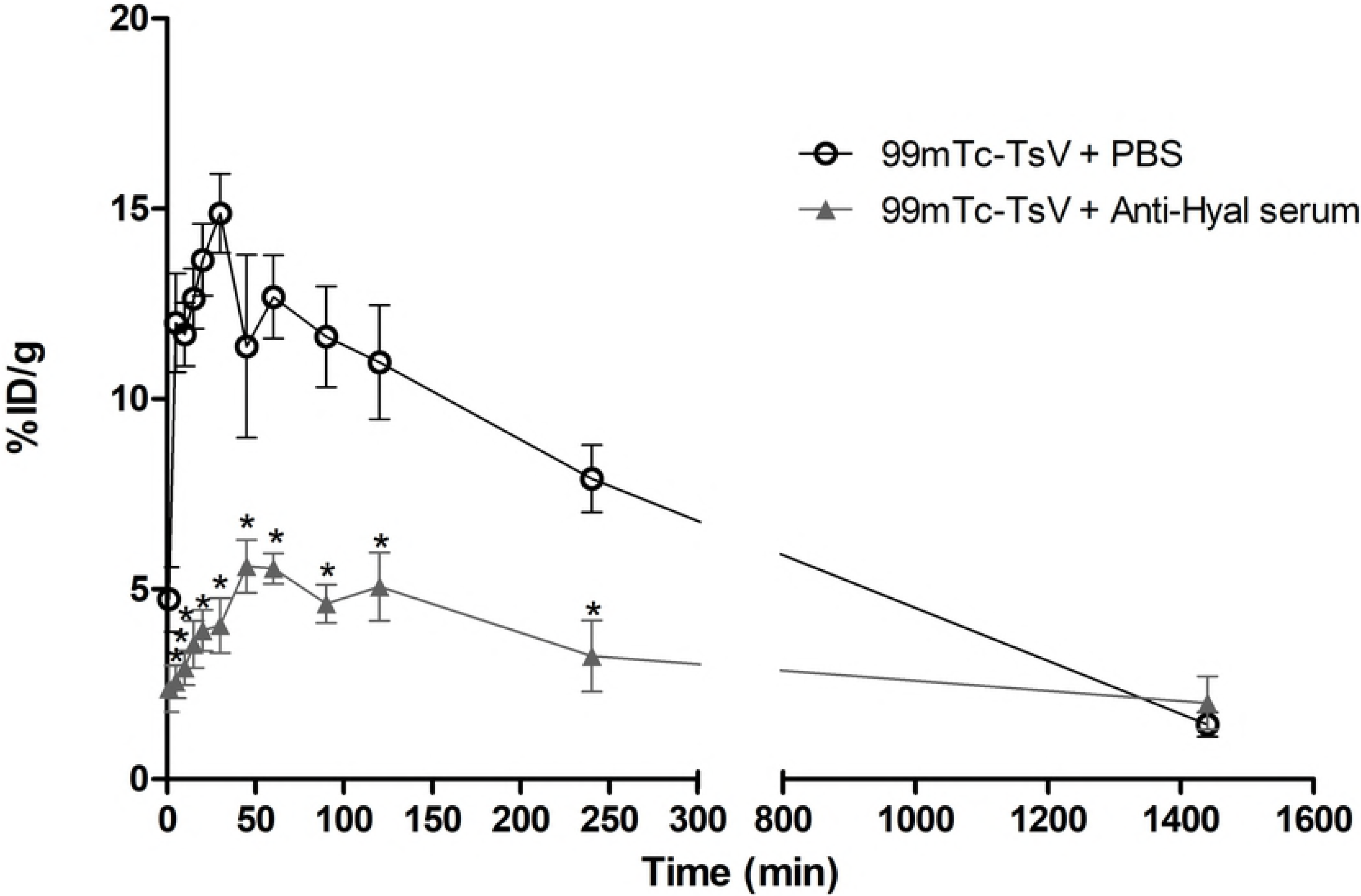
Blood clearance of ^99m^Tc-TsV. 3.7 MBq of ^99m^Tc-TsV diluted in PBS (^99m^Tc-TsV + PBS) or neutralized with anti-hyaluronidase serum (^99m^Tc-TsV + Anti-Hyal serum) was intramuscularly injected in Swiss mice (6-8 weeks old, 24-28 g; n = 6 per group). Radioactivity levels were measured in blood samples at 1, 5, 10, 15, 20, 30, 45, 60, 90, 120, 240, and 1440 minutes post-injection. Data are represented as the mean percentage of the injected dose of ^99m^Tc-TsV per gram of blood (% ID/g) ± S.E.M. of the mean. Values represent duplicates from two independent experiments. Statistical analyses were performed using a linear mixed model. Serum administration (p < 0.0001), time (p < 0.0001), and their interaction (p < 0.0001) had a statistically relevant effect on the mean ^99m^Tc-TsV blood clearance.

In contrast, the ^99m^Tc-TsV complex pre-neutralized with anti-hyaluronidase serum reaches lower levels in the bloodstream compared to the ^99m^Tc-TsV in PBS. This result shows that neutralization of TsV hyaluronidase activity significantly reduces TsV spreading from the injection site to the blood circulation.

Scintigraphic images of the mice corroborated the blood clearance results and showed that ^99m^Tc-TsV diluted in PBS quickly spreads from the injection site in the right tight muscle to the whole body between 30 and 120 min after TsV injection (Fig. 4A). On the other hand, when ^99m^Tc-TsV was neutralized with anti-hyaluronidase serum, TsV spreading from the injection site was visibly reduced at all times evaluated. Noteworthy, the labeled neutralized TsV remained at the site of injection (right tight muscle) (Fig. 4B).

**Figure 4.**
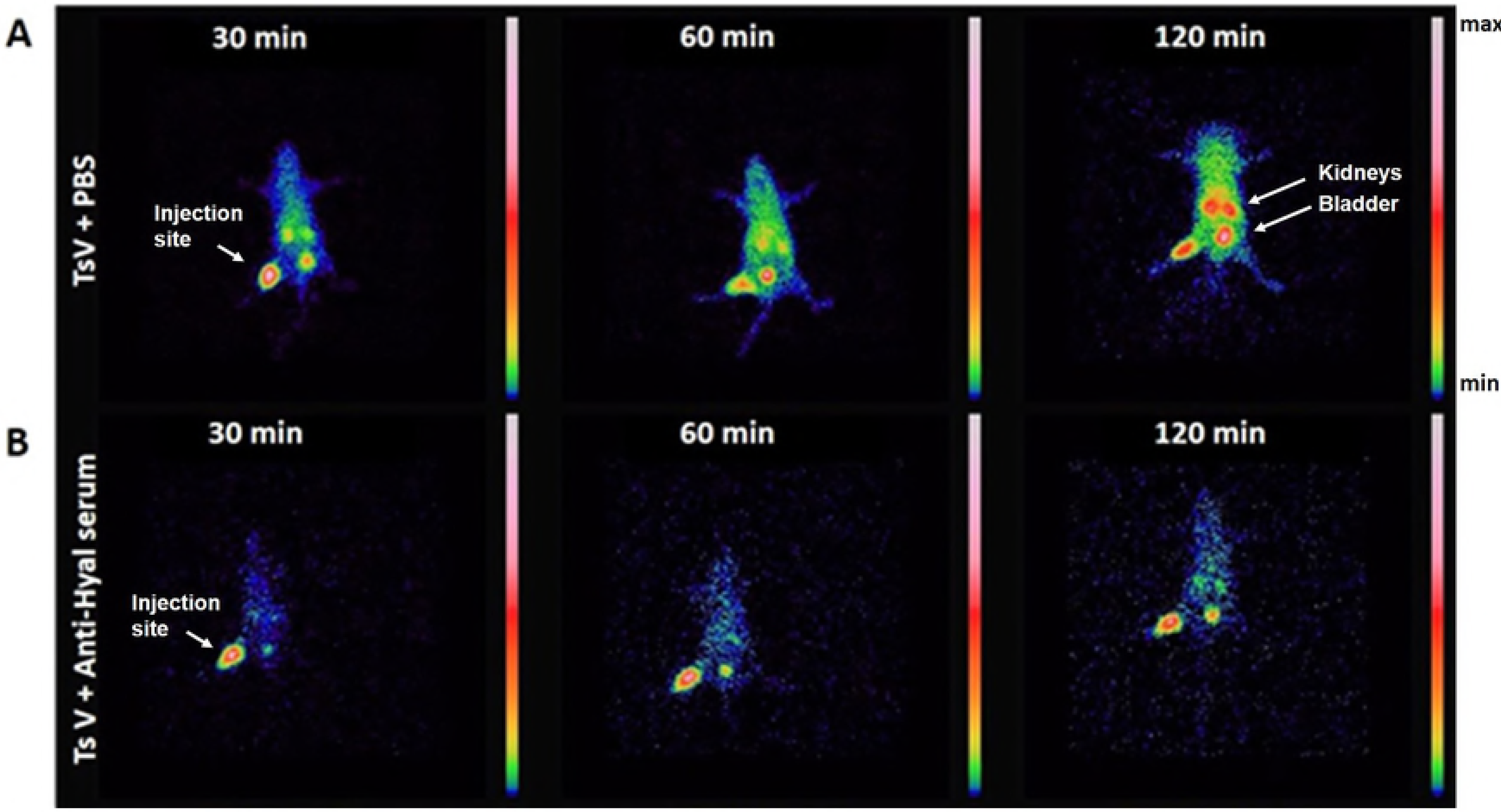
^99m^Tc-TsV spreading in mice over time. Representative scintigraphic images of mice injected with 18 MBq ^99m^Tc-TsV diluted in PBS (A) or neutralized with anti-hyaluronidase serum (B). Samples were intravenously injected in Swiss mice (6-8 weeks old, 24-28 g; n = 3 per group). Radioactivity levels were measured 30, 60, and 120 min post-injection. Images show a quick and growing spread of ^99m^Tc-TsV diluted in PBS over time (A). On the other hand, TsV neutralized with anti-hyaluronidase serum remains at the injection site (right tight muscle) (B). Images are pseudocolored according to the color scale.

Regarding the kinetics of TsV spreading, high levels of ^99m^Tc-TsV diluted in PBS were absorbed by the organs, particularly the kidneys and bladder (Fig. 4A), while lower levels of ^99m^Tc-TsV neutralized with anti-hyaluronidase serum reached these tissues over time (Fig. 4B).

### Neutralization of hyaluronidase delays biodistribution of TsV to tissues

Tissues such as heart, liver, lungs, spleen, and thyroid displayed different uptake levels of ^99m^Tc-TsV diluted in PBS and ^99m^Tc-TsV neutralized with anti-hyaluronidase serum (Fig. 5). Higher concentration of ^99m^Tc-TsV diluted in PBS was initially detected in these tissues (30 min), indicating a quick biodistribution of TsV from the injection site to the bloodstream and subsequently to the organs. After reaching the organs, labeled TsV concentration decreased over time, thus suggesting the elimination of the radiolabeled complex from the mice’s body. In agreement, ^99m^Tc-TsV levels increased in the kidneys, pointing towards renal elimination.

**Figure 5.**
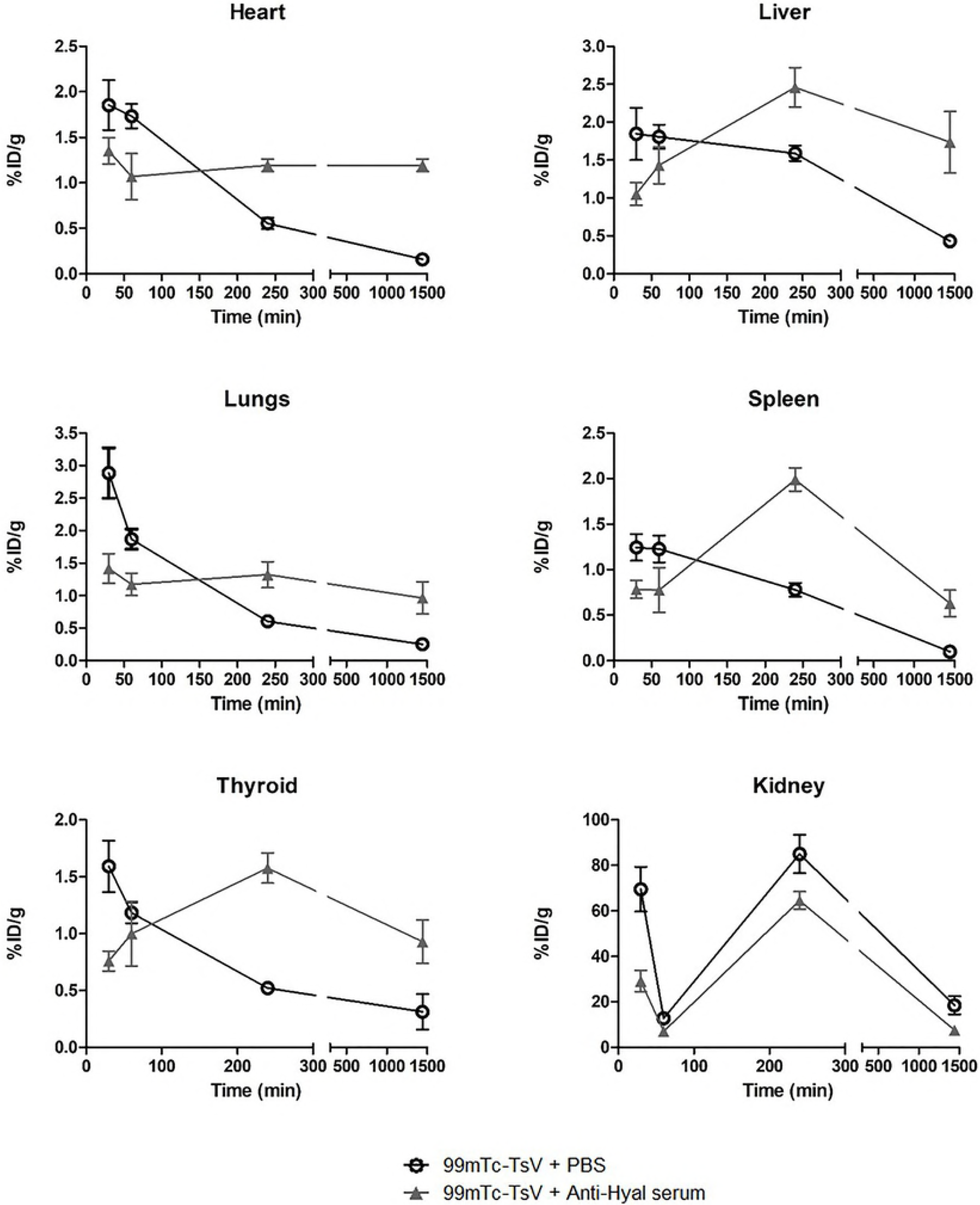
Biodistribution of ^99m^Tc-TsV. ^99m^Tc-TsV (3.7 MBq) diluted in PBS (^99m^Tc-TsV + PBS) or neutralized with anti-hyaluronidase serum (^99m^Tc-TsV + Anti-Hyal serum) was intramuscularly injected in Swiss mice (6-8 weeks old, 24-28 g; n = 6 per group). Radioactivity levels were measured in the heart, liver, lungs, spleen, thyroid and kidneys at 30, 60, 240 and 1440 min post-injection. The results are expressed as the percentage of injected dose/g of tissue (%ID/g). All values are presented as the mean ± S.E.M. of two independent experiments. Statistical analysis was performed using two-way ANOVA (factors: serum administration and time). Anti-hyaluronidase serum significantly affected the mean distribution of TsV to the liver (p < 0.0001), spleen (p = 0.0115), and kidneys (p = 0.0009), while time was a significant factor for TsV distribution to the heart (p < 0.0001), liver (p < 0.0001), lungs (p = 0.0095), spleen (p = 0.0008), and kidneys (p < 0.0001). A significant interaction between serum administration and time was observed in the heart (p = 0.00003), liver (p < 0.0001), spleen (p = 0.0337), and thyroid (p < 0.0001).

In contrast, the neutralized ^99m^Tc-TsV was initially detected at lower concentrations in the heart, liver, lungs, spleen, and thyroid, only reaching high levels at 240 min post-injection, after which time levels begin to decrease again (Fig. 5). Thus, hyaluronidase inhibition delays TsV incorporation into the bloodstream and organs. Moreover, lower concentrations of neutralized TsV reached the kidneys, when compared with ^99m^Tc-TsV diluted in PBS.

### TsV hyaluronidase neutralization as a first-aid treatment for scorpion sting

A blood clearance test was performed to evaluate the efficiency of anti-hyaluronidase serum to neutralize previously injected TsV *in situ*, thus simulating a first-aid treatment for scorpion sting.

Animals injected with ^99m^Tc-TsV diluted in PBS (control group) presented a quick absorption of TsV from the injection site into the bloodstream, followed by a decrease in blood concentration (Fig. 6), corroborating the results previously observed in the blood clearance assay (Fig. 3). On the other hand, animals treated with anti-hyaluronidase serum injected in the right tight muscle at 0, 10, and 30 min post-injection of ^99m^Tc-TsV diluted in PBS showed higher concentrations of labeled TsV in the bloodstream over time at all times tested (Fig. 6A-C). The increased concentration of neutralized TsV in the bloodstream indicates hindered biodistribution to the tissues (Fig. 6). Altogether, our data reveal the potential use of hyaluronidase inhibition as a novel first-aid strategy in envenoming.

**Figure 6.**
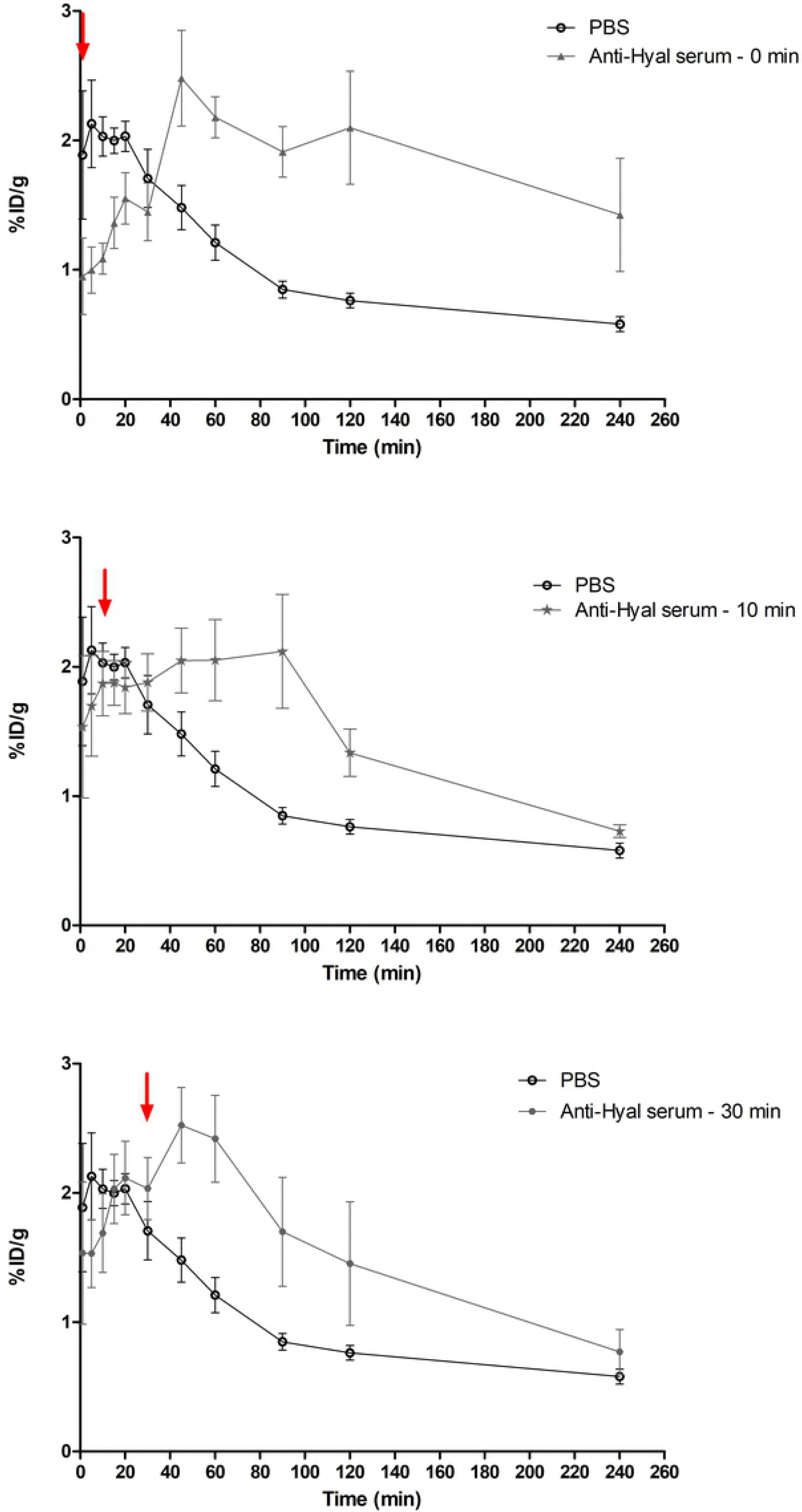
Hyaluronidase neutralization as a first-aid treatment for scorpion sting. 3.7 MBq of ^99m^Tc-TsV diluted in PBS was intramuscularly injected in Swiss mice (6-8 weeks old, 24-28 g; n=6 per group). Subsequently, anti-hyaluronidase serum was injected in the same site of ^99m^Tc-TsV injection at different times (Anti-Hyal serum; 0, 10, and 30 min post-injection; arrows indicate the injection times). Radioactivity levels in the bloodstream were measured at 1, 5, 10, 15, 20, 30, 45, 60, 90, 120, and 240 min post-injection of ^99m^Tc-TsV. All data are expressed as the mean percentage of the injected dose of ^99m^Tc-TsV per gram of blood (% ID/g) ± S.E.M. of the mean. Values are representative of duplicates from two independent experiments. Statistical analysis was performed using a linear mixed model. Time (p < 0.0001) and time x Anti-Hyal serum administration interaction (p < 0.0001) had a statistically relevant effect on the mean ^99m^Tc-TsV blood clearance.

## Discussion

Hyaluronidases are a class of enzymes produced by a great diversity of organisms, from bacteria to mammals [34,35]. They are involved in different processes such as inflammation, angiogenesis, embryogenesis, wound healing, tumor growth and progression, and systemic diffusion of venom toxins [36–39]. The role of hyaluronidases in venom spreading has been investigated, as this enzyme is a component vastly described both in vertebrate and invertebrate animal venoms. However, no study so far has shown how hyaluronidase activity interferes with venom distribution [17,40–44].

Previous studies have demonstrated the role of hyaluronidase in venom dissemination [36,44], in enhancing the effects of hemorrhagic toxins from snake venoms [27,37], in triggering allergic reactions to bee and wasp venoms [45–47], and in increasing the effects of other toxins from spider and scorpion venoms [14,48]. Moreover, Horta and collaborators [16] have greatly advanced the understanding of hyaluronidase in *T. serrulatus* through characterization studies showing the role of this enzyme in venom lethality. However, evidence demonstrating the role of hyaluronidases in venom spreading and describing the steps from scorpion sting to venom elimination was still lacking. In the present study, we demonstrated how the inhibition of TsV hyaluronidase activity affects venom biodistribution.

Our data show that TsV distribution kinetics is fast and efficient. TsV is distributed from the injection site to the bloodstream and organs in the first 30 min post-injection (Fig. 3, 4A and 5). After that time, the level of TsV decreases in the organs (Fig. 5) and increases in the kidneys (Fig. 4A, 5), thus indicating a renal route of elimination of the radiolabeled complex ^99m^Tc-TsV. This shows that the venom is quickly biodistributed from the injection site to the target organs, where it activates receptors and other biological targets. The binding triggers signaling cascades that culminate in all the symptomatology of scorpion poisoning. Following biodistribution, renal excretion is an important route of elimination of TsV from the organism [49,50].

Previous studies have used a toxic fraction isolated from *T. serrulatus* venom radiolabeled with 99m-Tc (^99m-Tc^TsTx) for biodistribution assays in young rats [49]. In that study, it was observed that the isolated fraction is first detected at low levels in the organs and reaches its maximum concentration in the brain, heart, thyroid, lungs, spleen, liver, and blood after 60 to 180 min. In the kidneys, the highest concentration of ^99m-Tc^TsTx is detected 360 min after injection [49]. The biodistribution and elimination of TsTx are slower compared to that of the total venom observed in the present study and may be explained by the low molecular weight of the TsTx fraction (~7 kDa) that lacks hyaluronidase (~ 50 kDa) in its composition.

Studies of venom biodistribution and neutralization of circulating venom allow a better understanding of the pathophysiology of envenomation, especially through the determination of venom levels in tissues [49,51–55].

In the present work, we observed that the inhibition of hyaluronidase in TsV delays the process of venom distribution. Higher levels of TsV are detected at the injection site, and reduced levels are detected in the organs at early times (30 min) when compared to the control group (Fig. 4B and 5). The immunoneutralized venom is retained in the right tight muscle, and its spread from the injection site to the bloodstream is reduced (Fig. 4B). Over time, the levels of TsV in the organs show delayed increase when compared to control. The highest concentrations of immunoneutralized venom are observed in the tissues 240 min after injection, which represents a 190-min delay in comparison with the untreated venom. Lower levels of immunoneutralized TsV were also observed in the kidneys, in relation to the control, indicating reduced renal clearance (Fig 4 and 5). The delay in the biodistribution of TsV caused by inhibition of hyaluronidase may compromise the synergistic action of the venom’s components, which are relevant in the envenoming process, and may culminate with the reduction of TsV lethality observed by Horta et al. [16].

Revelo et al. [51] demonstrated the effect of *T. serrulatus* antivenom on the biodistribution of TsV. High levels of venom were detected in mice serum and organs up to 8 h after subcutaneous injection of TsV. In contrast, when antivenom was applied intravenously at times 0 or 1 h after venom injection, venom levels detected in blood and tissues were significantly reduced [51]. These data show the effectiveness of antivenom in blocking venom biodistribution and indicate that anti-hyaluronidase antibodies may exist in total antivenom.

Due to the hyaluronidase action facilitating initial venom dissemination, we hypothesized that anti-hyaluronidase serum could complement anti-serum therapy as a first-aid treatment for envenomation. Some studies point to the use of hyaluronidase activity inhibition in envenoming processes as a first-aid action [56]. Here we proceeded with inhibition of hyaluronidase after venom injection at times 0, 10, and 30 min, and observed a higher concentration of labeled TsV in the bloodstream until 120 or 240 minutes after TsV injection, depending on anti-hyaluronidase administration time, when compared to the control group (Fig. 6). These results corroborated the ^99m^Tc-TsV biodistribution assays previously neutralized with anti-hyaluronidase serum, which showed that the maximum concentration of TsV detected in tissues occurs 240 min after injection (Fig. 5). Thus, aiming at using hyaluronidase neutralization as a first-aid treatment, our results were effective in showing a delay in the biodistribution of TsV to target organs and its accumulation in the bloodstream. In a real envenoming situation, delaying venom biodistribution may compensate for the time required for the sting victims to seek medical attention and treatment with antivenom serum, especially in remote locations with poor access to hospitals. This would represent a breakthrough in the treatment of systemic envenoming by venomous animals, which are considered neglected issues by the World Health Organization (WHO) due to the lack of adequate access to antivenom therapy where they are needed [57].

In addition, the accumulation of TsV in the bloodstream as a result of hyaluronidase activity neutralization (Fig. 6) suggests a correlation between hyaluronidase activity and venom flow from the bloodstream to the tissues. These results indicate that TsV hyaluronidase is important not only to allow venom access from the sting/bite site to the bloodstream (Fig. 3, 4) but is also involved in the biodistribution of TsV from the blood to the organs (Fig. 5, 6).

In endothelial cells, hyaluronic acid (HA) stimulates cell proliferation, migration, and neovascularization, and regulates endothelial barrier function [58]. As a key component of the glycocalyx in the vascular wall, HA is crucial for vascular integrity and maintenance of blood vessel continuity [59]. Especially in the glomerulus, HA is pivotal to the integrity of protein permeability barrier [60,61]. Our results suggest that TsV hyaluronidase is relevant to the cleavage of the HA present in the endothelial barrier and, therefore, promotes the biodistribution of TsV from the blood to the tissues.

Studies of this nature can contribute to the development of more effective envenomation therapies and help clarify the mechanisms of action of components from the TsV. Herein, hyaluronidase was shown as a key enzyme for the biodistribution of TsV from the venom injection site to the bloodstream and subsequently to the target tissues. This enzyme promotes the rapid distribution of TsV toxins through the victim's body and is pivotal in the envenoming process. Since we have proved the critical role of hyaluronidase in scorpionism, our findings lead the way for new therapeutic strategies.

## SUPPORTING INFORMATION CAPTIONS

All the data supporting the findings of this study are included in the article.

## References

1. Academia Brasileira de Ciências (ABC). 2010. Doençaas negligenciadas. Ciência e Tecnologia para o Desenvolvimento Nacional: Estudos Estratégicos./Souza W, coordinator. Rio de Janeiro: Academia Brasileira de Ciências. 56 p

2. Ministério da Saúde do Brasil. SINAN - Sistema de informação de agravos de notificação. 2018. Available from: http://portalsinan.saude.gov.br/o-sinan

3. Bucaretchi F, Baracat ECE, Nogueira RJN, Chaves A, Zambrone FAD, Fonseca MRCC, et al. A comparative study of severe scorpion envenomation in children caused by Tityus bahiensis ans Tityus serrulatus. Rev Inst Med Trop Sao Paulo. 1995;37(4):331–6.

4. Freire-Maia L, Campos JA, Amaral CFS. Approaches to the treatment of scorpion envenoming. Toxicon. 1994;32(9):1009–14.

5. Eickstedt, V. R. D.; Candido, D. M.; Jorge, M. T.; Albuquerque, M. J.; Ribeiro LA. Escorpionismo no Estado de São Paulo: ocorrência de Tityus serrulatus e T. bahiensis, frequência de acidentes e gravidade dos envenenamentos. Rev Inst Med Trop. 1994;27:56.

6. R LW, Lourenço WR. Parthenogenesis in Scorpions: Some History – New Data. J Venom Anim Toxins incl Trop Dis V J Venom Anim Toxins incl Trop Dis. 2008;14(1):19–44.

7. Pucca MB, Cerni FA, Lopes E, Junior P, Bordon KDCF, Amorim FG, et al. Tityus serrulatus venom – a lethal cocktail. Toxicon. 2015; 108:272–84.

8. Venancio EJ, Portaro FC V, Kuniyoshi AK, Carvalho DC, Pidde-Queiroz G, Tambourgi D V. Enzymatic properties of venoms from Brazilian scorpions of Tityus genus and the neutralisation potential of therapeutical antivenoms. Toxicon. 2013;69:180–90.

9. Cupo P, Jurca M, Azevedo-Marques MM, Oliveira JSM, Ilering SE. Severe scorpion envenomation in Brazil. Clinical, laboratory and anatomopathological aspects. Rev Inst Med Trop Sao Paulo. 1994;36(1):67–76.

10. Chippaux JP, Goyffon M. Epidemiology of scorpionismo: A global appraisal. Acta Trop. 2008;107(2):71–9.

11. Cupo P. Clinical update on scorpion envenoming. Rev Soc Bras Med Trop. 2015;48(6):642–9.

12. Alvarenga ER, Mendes TM, Magalhaes BF, Siqueira FF, Dantas AE, Barroca TM, et al. Transcriptome analysis of the Tityus serrulatus scorpion venom gland. Open J Genet. 2012;02(04):210–20.

13. Verano-Braga T, Rocha-Resende C, Silva DM, Ianzer D, Martin-Eauclaire MF, Bougis PE, et al. Tityus serrulatus Hypotensins: a new family of peptides from scorpion venom. Biochem Biophys Res Commun. 2008;371(3):515–20.

14. Pessini AC, Takao TT, Cavalheiro EC, Vichnewski W, Sampaio S V, Giglio JR, et al. A hyaluronidase from Tityus serrulatus scorpion venom : isolation, characterization and inhibition by ¯ avonoids. Toxicon. 2001;39:1495–504.

15. Carmo AO, Oliveira-Mendes BBR, Horta CCR, Magalhães BF, Dantas AE, Chaves LM, et al. Molecular and functional characterization of metalloserrulases, new metalloproteases from the Tityus serrulatus venom gland. Toxicon. 2014;90:45–55.

16. Horta CCR, Magalhães B de F, Oliveira-Mendes BBR, Carmo AO do, Duarte CG, Felicori LF, et al. Molecular, Immunological, and Biological Characterization of Tityus serrulatus Venom Hyaluronidase: New Insights into Its Role in Envenomation. PLoS Negl Trop Dis. 2014 Feb 13;8(2):e2693.

17. Bordon KCF, Wiezel GA, Amorim FG, Arantes EC. Arthropod venom Hyaluronidases: biochemical properties and potential applications in medicine and biotechnology. J Venom Anim Toxins Incl Trop Dis. 2015;21(1):43.

18. Cologna CT, Marcussi S, Giglio JR, Soares AM, Arantes EC. Tityus serrulatus Scorpion Venom and Toxins : An Overview. Protein Pept Lett. 2009;920–32.

19. Rates B, Ferraz KK, Borges MH, Richardson M, Lima ME De, Pimenta AMC. Tityus serrulatus peptidomics: acessing venom peptide diversity. Toxicon. 2008; 52(5):611–8.

20. Rodríguez de la Vega RC, Possani LD. Current views on scorpion toxins specific for K+-channels. Toxicon. 2004;43(8):865–75.

21. Rodríguez de la Vega RC, Possani LD. Overview of scorpion toxins specific for Na+ channels and related peptides: biodiversity, structure-function relationships and evolution. Toxicon. 2005;46(8):831–44.

22. Ferreira LAF, Alves EW, Henriques OB. Peptide T, a novel bradykinin potentiator isolated from Tityus Serrulatus scorpion venom. Toxicon. 1993;31(8):941–7.

23. Chavez-Olortegui C, Kalapothakis E, Ferreira AMBM, Ferreira AP, Diniz CR. Neutralizing capacity of antibodies elicited by a non-toxic protein purified from the venom of the scorpion Tityus serrulatus. Toxicon. 1997;35(2):213–21.

24. Harris F, Dennison SR, Phoenix DA. Anionic Antimicrobial Peptides from Eukaryotic Organisms. Curr protein Pept Sci. 2009;10:585–606.

25. Khan N, Niazi ZR, Rehman F ur, Akhtar A, Khan MM, Khan S, et al. Hyaluronidases: A Therapeutic Enzyme. Protein Pept Lett. 2018 Sep 6;25(7):663–76.

26. Duran-Reynals F. Studies On A Certain Spreading Factor Existing In Bacteria and its Significance For Bacterial Invasiveness. J Exp Med. 1933;58(2):161–81.

27. Tu AT, Hendon RR. Characterization of lizard venom hyaluronidase and evidence for its action as a spreading factor. Comp Biochem Physiol. 1983;76(2):377–83.

28. Possani DL, Alagón AC, Fletcher PL, Erickson B. Purification and Properties of Mammalian Toxins from the Venom of the Brazilian Scorpion Tityus serrulatus Lutz and Mello. Arch Biochem Biophys. 1977;180:394–403.

29. Amorim FG, Boldrini-frança J, Castro K De, Bordon F, Cardoso IA. Heterologous expression of rTsHyal-1 : the first recombinant hyaluronidase of scorpion venom produced in Pichia pastoris system. Appl Microbiol Biotechnol. 2018 Apr;102(7):3145–3158.

30. Lowry OH, Rosebrough NJ, Farr L, Randall RJ. Protein measurement with the Folin Phenol reagent. J Biol Chem. 1951;193(1):265–75.

31. Pukrittayakamee S, Warrell DA, Desakorn V, McMichael AJ, White NJ, Bunnag D. The hyaluronidase activities of some Southeast Asian snake venoms. Toxicon. 1988;26(7):629–37.

32. Fernandes RS, Silva J de O, Lopes SCA, Chondrogiannis S, Rubello D, Cardoso VN, et al. Technetium-99m-labeled doxorubicin as an imaging probe for murine breast tumor (4T1 cell line) identification. Nucl Med Commun. 2016 Mar;37(3):307–12.

33. USP 34. The United States Pharmacopoeia National Formulary.Rockville: United States Pharmacopoeial Convention Inc. 2012.

34. Stern R, Jedrzejas MJ. Hyaluronidases: Their Genomics, Structures, and Mechanisms of Action. Chem Rev. 2006 Mar;106(3):818–39.

35. Deangelis PL, Liu J, Linhardt RJ. Chemoenzymatic synthesis of glycosaminoglycans: Re-creating, re-modeling and re-designing nature’s longest or most complex carbohydrate chains. Glycobiology. 2013;23(7):764–77.

36. Girish KS, Kemparaju K. The magic glue hyaluronan and its eraser hyaluronidase: a biological overview. Life Sci 2007 May; 80(21):1921–43.

37. Kemparaju K, Girish KS. Snake venom hyaluronidase: a therapeutic target. Cell Biochem Funct. 2006;24(1):7–12.

38. Lokeshwar VB, Selzer MG. Hyaluronidase: both a tumor promoter and suppressor. Semin Cancer Biol. 2009;18(4):281–7.

39. Jiang D, Liang J, Noble P. Hyaluronan as an immune regulator in human diseases. Physiol Ver. 2011;91:221–64.

40. Barth T, Mandacaru SC, Charneau S, Souza MV de, Ricart CAO, Noronha EF, et al. Biochemical and structural characterization of a protein complex containing a hyaluronidase and a CRISP-like protein isolated from the venom of the spider Acanthoscurria natalensis. J Proteomics. 2018 Aug 27. pii: S1874-3919(18)30324–5.

41. Meyer K. Hyaluronidases. Enzym. 1971;5:307–20.

42. Poh CH, Yuen R, Chung MCM, Khoo HE. Purification and partial characterization of hyaluronidase from stonefish (Synanceja horrida) venom. Comp Biochem Physiol -- Part B Biochem. 1992;101(1–2):159–63.

43. Kreil G. Hyaluronidases--a group of neglected enzymes. Protein Sci. 1995 Sep;4(9):1666–9.

44. Fox JW. A brief review of the scientific history of several lesser-known snake venom proteins: l-amino acid oxidases, hyaluronidases and phosphodiesterases. Toxicon. 2013 Feb;62:75–82.

45. Rungsa P, Incamnoi P, Sukprasert S, Uawonggul N, Klaynongsruang S, Daduang J, et al. Cloning, structural modelling and characterization of VesT2s, a wasp venom hyaluronidase (HAase) from Vespa tropica. J Venom Anim Toxins Incl Trop Dis. 2016;22:28.

46. King TP, Wittkowski KM. Hyaluronidase and hyaluronan in insect venom allergy. Int Arch Allergy Immunol. 2011;156(2):205–11.

47. An S, Chen L, Wei JF, Yang X, Ma D, Xu X, et al. Purification and characterization of two new allergens from the venom of Vespa magnifica. PLoS One. 2012;7(2):5–12.

48. Ferrer VP, de Mari TL, Gremski LH, Trevisan Silva D, da Silveira RB, Gremski W, et al. A novel hyaluronidase from brown spider (Loxosceles intermedia) venom (Dietrich’s Hyaluronidase): from cloning to functional characterization. PLoS Negl Trop Dis. 2013;7(5):e2206.

49. Nunan EA, Moraes MF, Cardoso VN, Moraes-Santos T. Effect of age on body distribution of Tityustoxin from Tityus serrulatus scorpion venom in rats. Life Sci. 2003;73(3):319–25.

50. Oliveira-mendes BBR De, Campolina C, Horta R, Leal HG, Brandão-dias PFP, Eustáquio S, et al. CPP-Ts : a new intracellular calcium channel modulator and a promising tool for drug delivery in cancer cells. Sci Rep 2018;(September):1–13.

51. Revelo MP, Bambirra EA, Ferreira AP, Diniz CR, Chavez-Olortegui C. Body distribution of Tityus serrulatus scorpion venom in mice and effects of scorpion antivenom.. Toxicon. 1996;34(10):1119–25.

52. Ismail M, Abdullah ME, Morad AM, Ageel AM. Pharmacokinetics of 125I-labelled venom from the scorpion Androctonus amoreuxi, Aud. and Sav. Toxicon. 1980;18(3):301–8.

53. Ismail M, Shibl AM, Morad AM, Abdullah ME. Pharmacokinetics of125I-labelled antivenin to the venom from the scorpion Androctonus amoreuxi. Toxicon. 1983;21(1):47–56.

54. Ismail M, Abd-Elsalam MA. Are the toxicological effects of scorpion envenomation related to tissue venom concentration? Toxicon. 1988;26(3):233–56.

55. Dasgupta SC, Gomes A, Lahiri SC. Detection and distribution of scorpion (Heterometrus bengalensis) venom in rabbit tissues by enzyme-linked immunosorbent assay (ELISA). Indian J Exp Biol. 1991;29(7):691–2.

56. Girish KS, Kemparaju K. Inhibition of Naja naja venom hyaluronidase by plant-derived bioactive components and polysaccharides. Biochem. 2005;70(8):948–52.

57. World Health Organization (WHO). Rabies and envenoming: a neglected public health issue: report of a Consultative Meeting. Geneva: World Health Organization.2007: 21–22.

58. Genasetti A, Vigetti D, Viola M, Karousou E, Moretto P, Rizzi M, et al. Hyaluronan and human endothelial cell behavior. Connect Tissue Res. 2008;49(3–4):120–3.

59. Singleton PA. Hyaluronan Regulation of Endothelial Barrier Function in Cancer. Adv Cancer Res. 2014;91(2):191–209.

60. Dane MJC, Van Den Berg BM, Avramut MC, Faas FGA, Van Der Vlag J, Rops ALWMM, et al. Glomerular endothelial surface layer acts as a barrier against albumin filtration. Am J Pathol. 2013;182(5):1532–40.

61. Meuwese MC, Broekhuizen LN, Kuikhoven M, Heeneman S, Lutgens E, Gijbels MJJ, et al. Endothelial surface layer degradation by chronic hyaluronidase infusion induces proteinuria in apolipoprotein E-deficient mice. PLoS One. 2010;5(12):1–7.

